# The Distal Polybasic Cleavage Sites of SARS-CoV-2 Spike Protein Enhance Spike Protein-ACE2 Binding

**DOI:** 10.1101/2020.06.09.142877

**Authors:** Baofu Qiao, Monica Olvera de la Cruz

## Abstract

The receptor-binding domain (RBD) of the SARS-CoV-2 spike protein plays a crucial role in binding the human cell receptor ACE2 that is required for viral entry. Many studies have been conducted to target the structures of RBD-ACE2 binding and to design RBD-targeting vaccines and drugs. Nevertheless, mutations distal from the SARS-CoV-2 RBD also impact its transmissibility and antibody can target non-RBD regions, suggesting the incomplete role of the RBD region in the spike protein-ACE2 binding. Here, in order to elucidate distant binding mechanisms, we analyze complexes of ACE2 with the wild type spike protein and with key mutants via large-scale all-atom explicit solvent molecular dynamics simulations. We find that though distributed approximately 10 nm away from the RBD, the SARS-CoV-2 polybasic cleavage sites enhance, via electrostatic interactions and hydration, the RBD-ACE2 binding affinity. A negatively charged tetrapeptide (GluGluLeuGlu) is then designed to neutralize the positively charged arginine on the polybasic cleavage sites. We find that the tetrapeptide GluGluLeuGlu binds to one of the three polybasic cleavage sites of the SARS-CoV-2 spike protein lessening by 34% the RBD-ACE2 binding strength. This significant binding energy reduction demonstrates the feasibility to neutralize RBD-ACE2 binding by targeting this specific polybasic cleavage site. Our work enhances understanding of the binding mechanism of SARS-CoV-2 to ACE2, which may aid the design of therapeutics for COVID-19 infection.

TOC:
The SARS-CoV-2 spike protein-ACE2 complex showing the polybasic cleavage sites

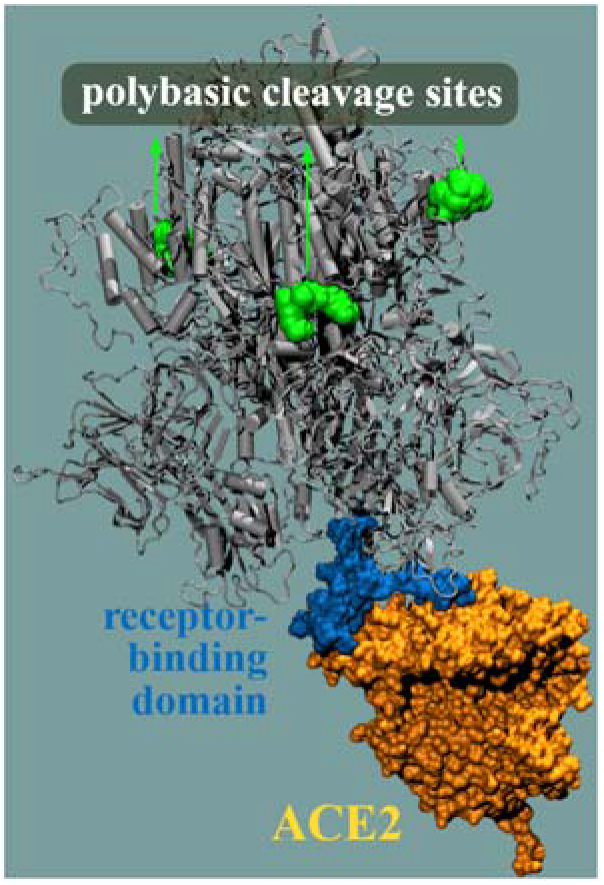

---

The outbreak of the SARS-CoV-2 related disease COVID-19 has caused more than 6 million infections and over 0.3 million deaths globally. The SARS-CoV-2 spike protein exhibits two notable features; its receptor-binding domain (RBD) is optimized to bind the human cell receptor angiotensin-converting enzyme 2 (ACE2) and each subunit of the spike protein trimer has a polybasic cleavage site. ^1^ Given the structural similarity between the spike proteins of the SARS-CoV-2 and its close relative SARS-CoV (2003), the RBD of the SARS-CoV-2 has been quickly recognized in targeting the ACE2 receptor, ^2–4^ which is required for subsequent viral entry. The atomic structures of the contact region between the SARS-CoV-2 RBD and ACE2 have been experimentally obtained by different groups. ^5–7^ Owing to the crucial role of the SARS-CoV-2 RBD, numerous vaccine and drug candidates have been proposed to target the RBD to inhibit the RBD-ACE2 binding. ^8–10^

However, more research is beginning to show that mutations distal from the RBD (non-RBD mutations) of SARS-CoV-2 also influence its transmissibility. For instance, Walls et al. ^2^ found that by mutating the T_678_NSPRRAR_685_ residues, which are distributed around 10 nm from the SARS-CoV-2 RBD, to a variant with T_678_--IL--R_685_, the transduction efficiency decreased in human ACE2-expressing Baby Hamster Kidney cells. The mutant D614G (a single change in the genetic code; D = aspartic acid, G = glycine) of the SARS-CoV-2 spike protein, which began spreading in Europe in February, displayed stronger transmissibility. ^11^ The D614 and G614 residues are located about 7 - 10 nm from the SARS-CoV-2 RBD. Surprisingly, Yuan et al. ^12^ found that the antibody CR3022 isolated from a recovered SARS-CoV patient targeted a highly conserved epitope of SARS-CoV-2 and SARS-CoV, which is also distal from their RBDs. That is, the antibody CR3022 does not directly bind the spike protein RBD. All these works support the significance of non-RBD mutations and non-RBD-targeting in vaccine and drug design, which remains elusive thus far.

In the present work, large scale all-atom explicit solvent molecular dynamics (MD) simulations reveal that the spike protein polybasic cleavage sites, which are distributed approximately 10 nm away from the RBD, can enhance the binding affinity between the SARS-CoV-2 RBD and ACE2. This information is used to design a negatively charged tetrapeptide, GluGluLeuGlu, that targets the polybasic cleavage sites. The negatively charged glutamic acids (Glu) are introduced to charge neutralize the polybasic cleavage sites and the nonpolar leucine (Leu) is included to increase the hydrophobicity of the tetrapeptide, which favors binding to the polybasic cleavage site. The tetrapeptide GluGluLeuGlu is found to bind to one specific polybasic cleavage site, resulting in a remarkable weakening (34%) of the spike protein-ACE2 binding energy. Our work provides guidelines to design therapeutic peptides to inhibit SARS-CoV-2 RBD-ACE2 binding.

## RESULTS AND DISCUSSION

### Impact of the Polybasic Cleavage Site on RBD-ACE2 Binding Affinity

Each subunit of the SARS-CoV-2 trimeric spike protein has one polybasic cleavage site (R_682_RAR_685_). ^1^ The polybasic cleavage sites (R_682_RAR_685_) have not been reported for SARS-CoV or any other lineage B coronaviruses, and therefore, they are thought to be unique to SARS-CoV-2. ^2, 13, 14^ In comparison to the extensive studies carried out on the SARS-CoV-2 RBD, the polybasic cleavage sites have thus far not been widely investigated, and their function remains elusive. They are believed to be related to the viral transmissibility of SARS-CoV-2. ^2, 14^ Moreover, they have been found to be essential for spike protein-driven viral entry into lung cells ^15^.

To elucidate the influence of the polybasic cleavage sites, the wild type (WT) spike protein-ACE2 complex was investigated along with two mutants (**Figure 1**). The SARS-CoV-2 spike protein-ACE2 complex was downloaded from the Zhang-Server, ^16^ which contains the full length of the wild type SARS-CoV-2 spike protein bound to the ACE2 receptor. The spike protein-ACE2 complex was solvated in a water box with a length of 16×18×24 nm^3^, where 0.15 M NaCl was included. All-atom MD simulation was conducted to equilibrate the system using the CHARMM 36m force field. ^17^ To preserve the spike protein-ACE2 binding structure the non-hydrogen atoms of the spike protein trimer and ACE2 were restrained, which were removed in the following simulations.

**Figure 1.**
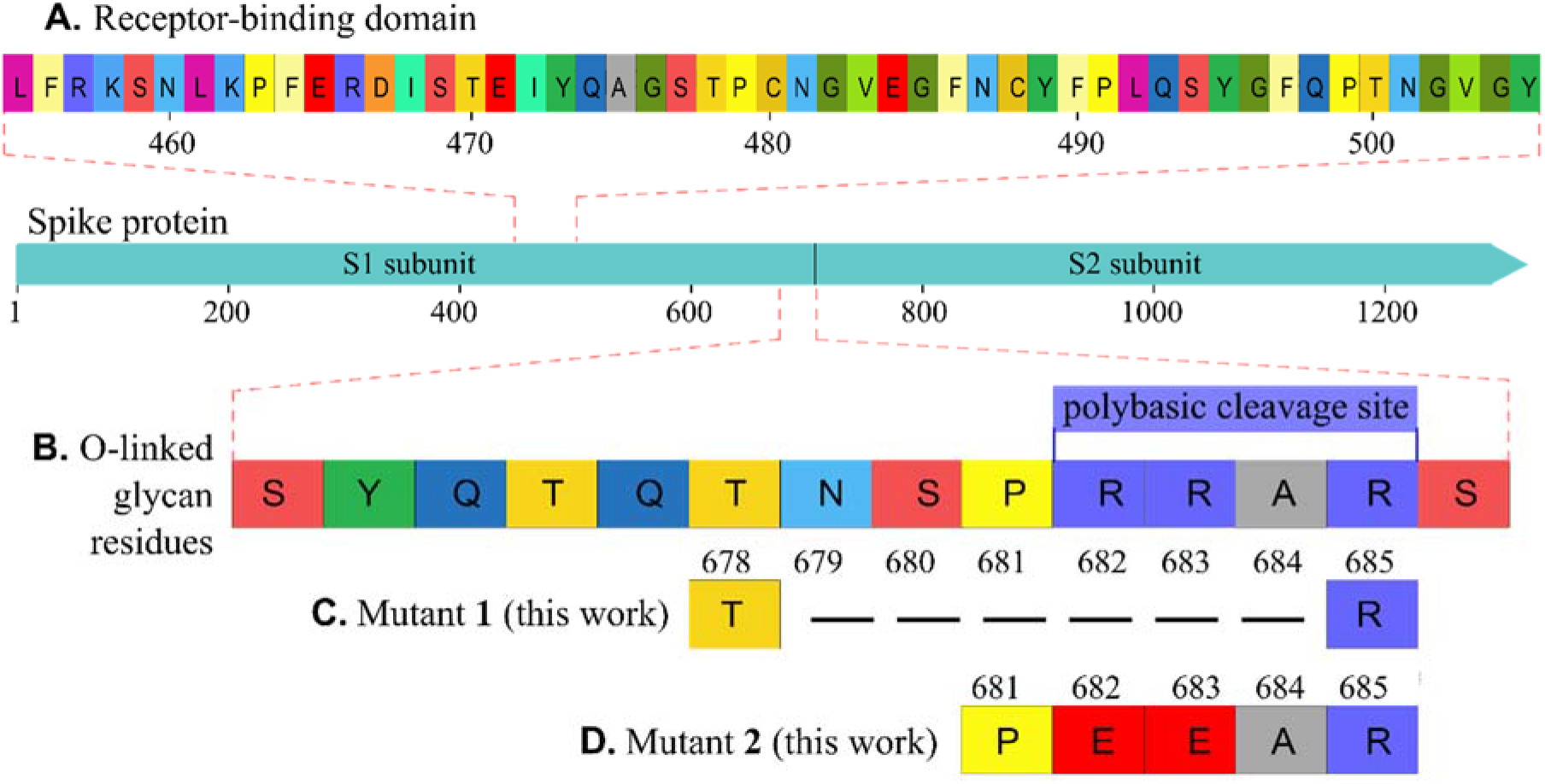
Wild type SARS-CoV-2 spike protein and the two mutants investigated here. (**A**) Receptor binding domain from L_456_ to Y_505_. ^1^ (**B**) O-linked glycan residues from S_673_ to S_686_, among which P_681_RRA_684_ is unique to SARS-CoV-2 compared to the other lineage B coronavirus. ^2, 13, 14^ R_682_RAR_685_ form a polybasic cleavage site. (**C**) Mutant **1,** where N_679_SPRRA_684_ residues were removed, is similar to the experimental variant ^2^ and (**D**) mutant **2** (E682E683), both investigated in this work.

The WT spike protein was subsequently mutated. For mutant **1**, the N_679_SPRRA_684_ residues were removed for all the three subunits of the spike protein trimer. This mutant is similar to the variant experimentally investigated (T_678_--IL--R_685_) ^2^. In the case of mutant **2**, the R_682_R_683_ residues were changed to E682E683 for all three subunits of the trimeric spike protein, where E stands for negatively charged glutamic acid. For each of the three systems (WT and mutants **1** and **2**) all-atom explicit solvent simulations were performed for a duration of 100 ns. And five parallel runs were carried out based on the simulation configurations at 20, 40, 60, 80, and 100 ns. See the Methods section. The five parallel simulations were employed to calculate the binding energy (Table S2 in the Supporting Information) and intermolecular hydrogen bonds between the SARS-CoV-2 RBD and ACE2.

Demonstrated in **Figure 2A** is the final simulation snapshot of the wild type spike protein-ACE2 complex. The polybasic cleavage sites are located 10 - 13 nm away from the RBD. In all three systems, the SARS-CoV-2 RBD stayed bound to ACE2 (**Figure 2B-D**). Nevertheless, the potential energy between the RBD and ACE2, which was the summation of the intermolecular short-range Coulomb and Lennard-Jones interactions, exhibited remarkable differences (**Figure 2E**). The RBD-ACE2 binding energy for the WT spike protein is −740 ± 70 kJ/mol. The removal of N_679_SPRRA_684_ compromises the RBD-ACE2 interaction by 36% to −470 ± 50 kJ/mol, supporting that mutant **1** heavily depresses the binding affinity between the RBD and ACE2. This is in line with the experimental finding that the mutation of T_678_NSPRRAR_685_ to a variant with T_678_--IL--R_685_ of a murine leukemia virus decreased the transduction efficiency of human ACE2-expressing Baby Hamster Kidney cells. ^2^ Meanwhile, the mutation from R_682_R_683_ to E_682_E_683_ decreased the RBD-ACE2 binding energy by 20% to −590 ± 40 kJ/mol. Therefore, both mutants greatly destabilized the RBD-ACE2 binding in comparison to the WT spike protein.

**Figure 2.**
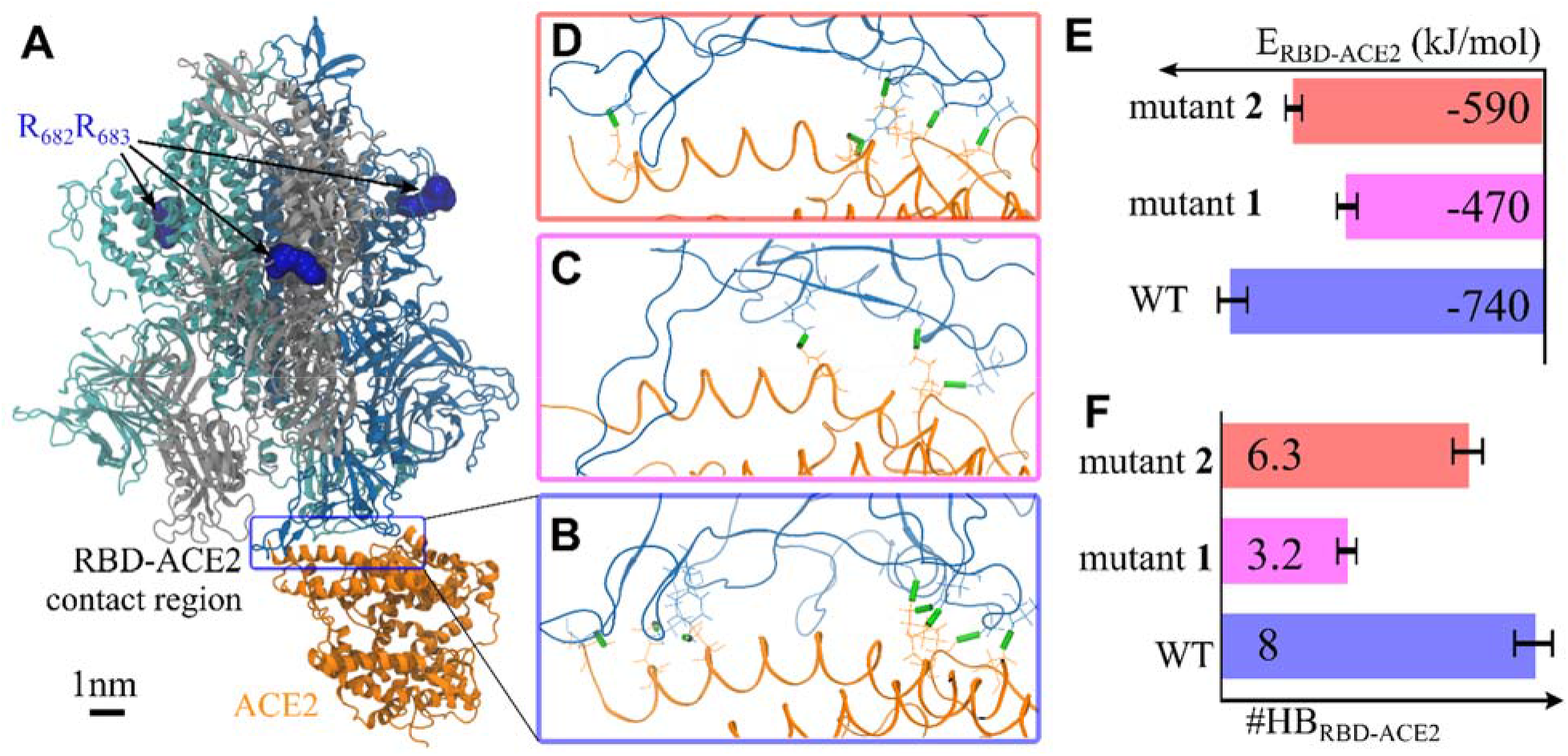
Distal mutants 1 and 2 both weaken the binding affinity between the RBD and ACE2. (**A**) Final simulation snapshot of the WT spike protein (in tan/gray/dark blue) and ACE2 (in orange). The R_682_R_683_ residues (blue beads) on the polybasic cleavage sites are approximately 10 - 13 nm away from the RBD. Water and salt ions are omitted for display. (**B**) The contact region between the WT spike protein (in dark blue) and ACE2 (in orange), where the RBD-ACE2 intermolecular hydrogen bonds are highlighted by green sticks. (**C**, **D**) are similar to (**B**), but for the mutants **1** and **2**, respectively. (**E**) The potential energy between the SARS-CoV-2 RBD and ACE2. See Table S2 for detailed values. (**F**) The number of intermolecular hydrogen bond (HB) between the RBD and ACE2. In (E, F), the averages and standard deviations are from five parallel runs (**Figure 4**).

Note that ACE2 is highly negatively charged (−28 *e*), which leads to a highly negatively charged extracellular membrane surface. The Coulomb interactions favor the adsorption of positively charged species, for instance, arginine residues. Also of note is that due to the positively charged nature, arginine has been found to enhance the cellular uptake of peptides ^18^ and promote the therapeutic delivery of peptides across the blood-brain barrier for Alzheimer’s disease ^19^. Therefore, the observed drops in the RBD-ACE2 binding energy for mutants **1** and **2** could be primarily ascribed to a change in (long-range) Coulomb interactions between ACE2 and the spike protein (Table S2), even though the polybasic cleavage sites are around 10 nm away from the RBD and ACE2. Moreover, the substitution of positively charged arginine with negatively charged glutamic acid increases protein hydration (Table S2), which is consistent with the fact that negatively charged amino acids exhibit stronger hydration than positively charged ones. ^20–22^

Further analysis of the intermolecular hydrogen bonds between the RBD and ACE2 demonstrated a similar influence of the mutants (**Figure 2F**). For the WT spike protein, there exist 8 ± 1 hydrogen bonds between the RBD and ACE2. This is in good agreement with the experimental finding ^5^ that eight RBD-ACE2 intermolecular polar interactions were suggested based on the cryo-EM structure. For mutant **1** the number of RBD-ACE2 intermolecular hydrogen bonds decreased by 60% to 3.2 ± 0.5, and by 21% (to 6.3 ± 0.8) for mutant **2**. These results further support that both mutants weaken the RBD-ACE2 binding affinity, in line with the calculated RBD-ACE2 intermolecular interaction energy.

Visualization of the distribution of the RBD-ACE2 intermolecular hydrogen bonds (**Figure 2B-D**) suggested that balanced hydrogen bonds at both ends of the RBD-ACE2 contact region play a crucial role in stabilizing the RBD-ACE2 binding. Specifically, the number of the intermolecular hydrogen bonds decreases at both termini from the WT spike protein to the mutant **2** and to the mutant **1**.

### Design of Oligopeptide Inhibitor

Interestingly, when we were preparing this work, an antibody CR3022 ^12^ was reported to bind a non-RBD region of the spike protein. Here, we explore the possibility to inhibit SARS-CoV-2 RBD-ACE2 binding by neutralizing the polybasic cleavage sites. To this end, we designed an oligopeptide: a tetrapeptide GluGluLeuGlu (EELE). Negatively charged glutamic acids (Glu) were included to neutralize the positively charged arginine residues at the polybasic cleavage sites. The hydrophobic leucine (Leu) residue was introduced to decrease the hydration, as a test simulation on a tetrapeptide EEEE showed that EEEE became quickly dissolved (within 10 ns) owing to the high hydration of the negatively charged amino acids. ^20–22^

Three EELE molecules were initially put next to the three polybasic sites of the trimeric spike protein. The system was equilibrated using the same process as that in the simulations in the absence of the tetrapeptide. One typical configuration is presented in **Figure 3A**. The inclusion of the tetrapeptide EELE can still preserve the spike protein-ACE2 complex (**Figure 3A, B**). Nevertheless, the RBD-ACE2 intermolecular potential energy (**Figure 3C**) remarkably dropped by 34% from −740 ± 70 kJ/mol without EELE to −490 ± 50 kJ/mol in the presence of EELE. Similarly, the number of RBD-ACE2 intermolecular hydrogen bonds dropped by 41%. Therefore, the presence of EELE greatly lessened RBD-ACE2 binding. Of note is that only one subunit of the trimeric spike protein directly binds to the ACE2 receptor. Consequently, the three polybasic cleavage sites have different distances to the ACE2 receptor and distinct local environment. Furthermore, the ACE2 receptor and the tetrapeptide EELE are both negatively charged (−28*e* for ACE2 and −3*e* for EELE). These effects collectively lead to the observation that the three tetrapeptides exhibited different binding behavior to the neighboring polybasic cleavage sites. Specifically, the polybasic cleavage site distributed the farthest from ACE2 stably binds to the tetrapeptide EELE for the whole simulation duration of 100 ns (**Figure 3A**); in contrast, the two polybasic cleavage sites closer to ACE2 form weaker interactions with their neighboring tetrapeptides EELE, which became unbound at around 40 ns and 74 ns.

**Figure 3.**
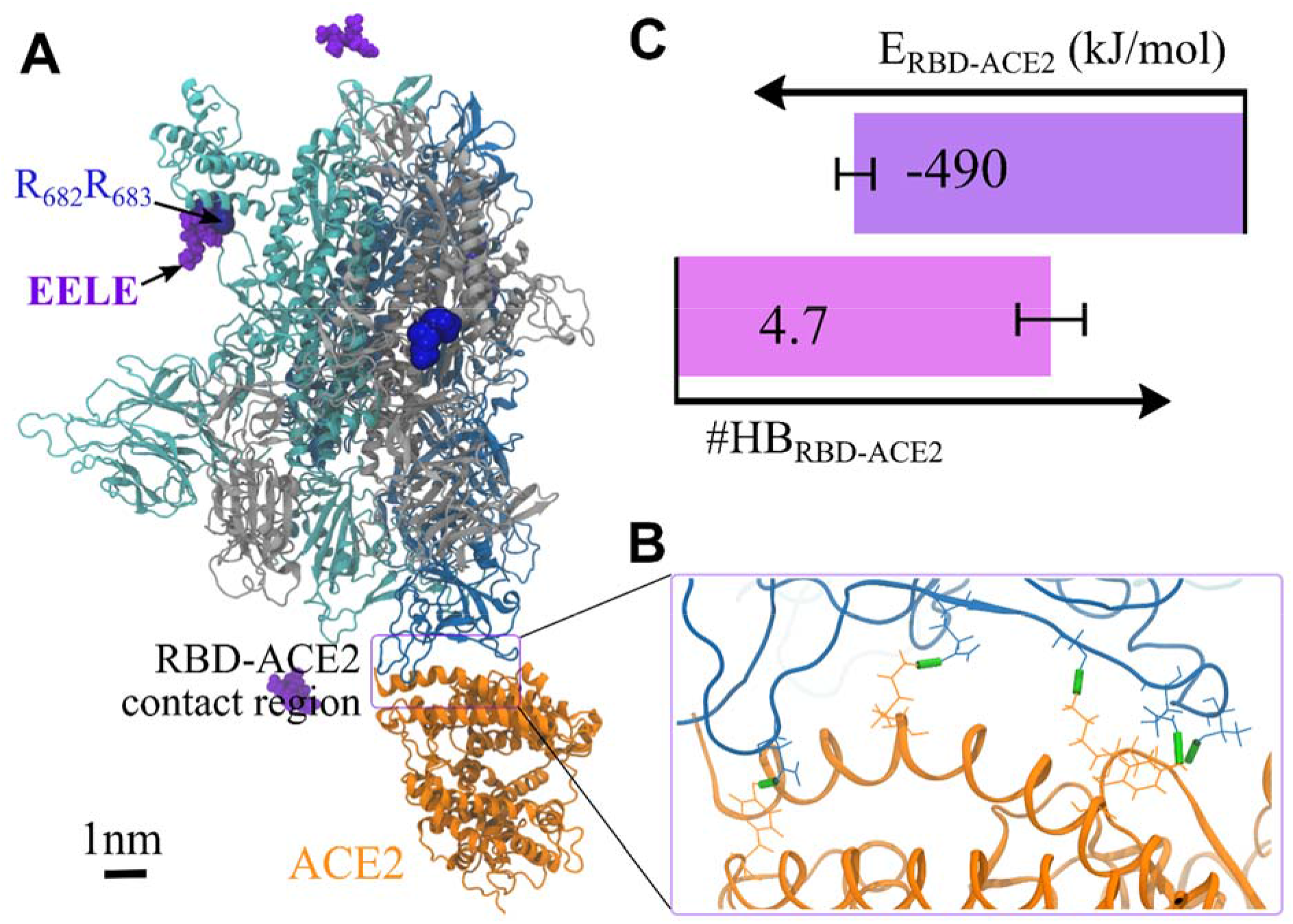
The presence of tetrapeptide EELE lessens the RBD-ACE2 binding. (**A**) Simulation snapshot of a spike-ACE2 complex in the presence of EELE (in violent). (**B**) The RBD-ACE2 contact region, where the RBD-ACE2 intermolecular hydrogen bonds are highlighted via green sticks. (**C**) Potential energy and number of hydrogen bonds between the RBD and ACE2. In (C), the averages and standard deviations are from five parallel runs (**Figure 4**).

To experimentally design polybasic cleavage site-targeting clinical therapeutic peptides, a further increase in the oligopeptide hydrophobicity is required to elevate the stability in the SARS-CoV-2 spike protein-oligopeptide binding to determine if binding to all of the polybasic cleavage sites reduces the RBD-ACE2 overall binding energy even further. Meanwhile, the delivery of therapeutic peptides is known to be challenging concerning their short half-life due to the rapid proteolytic degradation and short circulation time due to the low molecular weight. ^23, 24^ These could be potentially overcome by covalently grafting oligopeptides at the side chains of peptide brush polymers, ^25^ or integrating them onto peptide amphiphiles ^26^ or PEGylation ^27^.

## CONCLUSIONS

Using large-scale all-atom explicit solvent simulations, we investigated the impact of the SARS-CoV-2 polybasic cleavage sites, which are distributed approximately 10 nm away from the RBD, on the binding affinity of the RBD and ACE2. It is found that in comparison to the wild type SARS-CoV-2 spike protein, a mutant with the deletion of N_679_SPRRA_684_ and a mutant with the substitution of R_682_R_683_ to E_682_E_683_ can both lessen the binding strength of the SARS-CoV-2 RBD and ACE2. The mutation-driven difference is ascribed to the electrostatic interactions between the spike proteins (wild type and mutants) and ACE2 and their hydration. In line with recent experimental findings, ^2, 11^ this work supports that distal mutations can impact the SARS-CoV-2 RBD-ACE2 binding affinity. Our design of a tetrapeptide, GluGluLeuGlu, that binds to the polybasic cleavage site of SARS-CoV-2, demonstrates that the polybasic cleavage site is a target for neutralizing SAR-CoV-2 RBD-ACE2 binding. This supports a recent finding that the non-RBD region of SARS-CoV-2 can be targetted by an antibody. ^12^ This work besides shedding light on the mechanism by which the SARS-CoV-2 spike protein binds to human cells, suggests therapeutic peptides design to target the polybasic cleavage sites that inhibit SARS-CoV-2 RBD binding to ACE2.

## METHODS

### All-atom Simulations on the Wild Type SARS-CoV-2 Spike Protein–ACE2 Complex

All-atom explicit solvent molecular dynamics (MD) simulations were performed using the package GROMACS (version 2016.4). ^28^ The CHARMM 36m potential ^17^ was employed for the proteins, Na^+^ and Cl^−^ ions. The recommended CHARMM TIP3P water model ^29^ was employed with the structures constrained using the SETTLE algorithm ^30^.

The SARS-CoV-2 spike protein-ACE2 binding structure was downloaded from the Zhang-Server. ^16^ The spike protein-ACE2 complex was reconstructed using the C-I-TASSER model ^31^ based on the protein identification number QHD43416 ^32^ for the spike protein. Each subunit of the trimeric spike protein included the residues from M1 to T1273 with the net charge of −7*e*, and the ACE2 has the residues from S19 to D615 with the net charge of −28*e*, with *e* being the elementary charge.

The spike protein-ACE2 complex was first solvated in a simulation box with an edge length of 16×18×24 nm^3^ (**Figure 4**). 0.15 M NaCl was added along with 49 Na^+^ counterions of spike protein and ACE2. The system composition is listed in Table S1 in the Supporting Information. The energy of the simulation box was first minimized, which was followed by a simulation of 1 ps using the canonical ensemble (constant number of particles, volume, and temperature, NVT). The integration time step of 2 fs was employed with all the hydrogen-involved covalent bonds constrained using the LINCS algorithm ^33, 34^. Another simulation of 1 ps was subsequently conducted using the isothermal-isobaric ensemble (constant number of particles, pressure, and temperature, NPT). The velocity rescale temperature coupling was employed along with the Berendsen pressure coupling. In the following equilibration simulation of 10 ns, the Nosé-Hoover temperature coupling was applied along with the Parrinello-Rahman pressure coupling.^35^ In all the equilibration simulations above, the coordinates of the non-hydrogen atoms of both the spike protein trimer and ACE2 were restrained using a force constant of 1000 kJ/mol/nm^2^ to preserve the spike protein-ACE2 binding structure. The other parameters were the same as those in the production simulation below.

**Figure 4.**
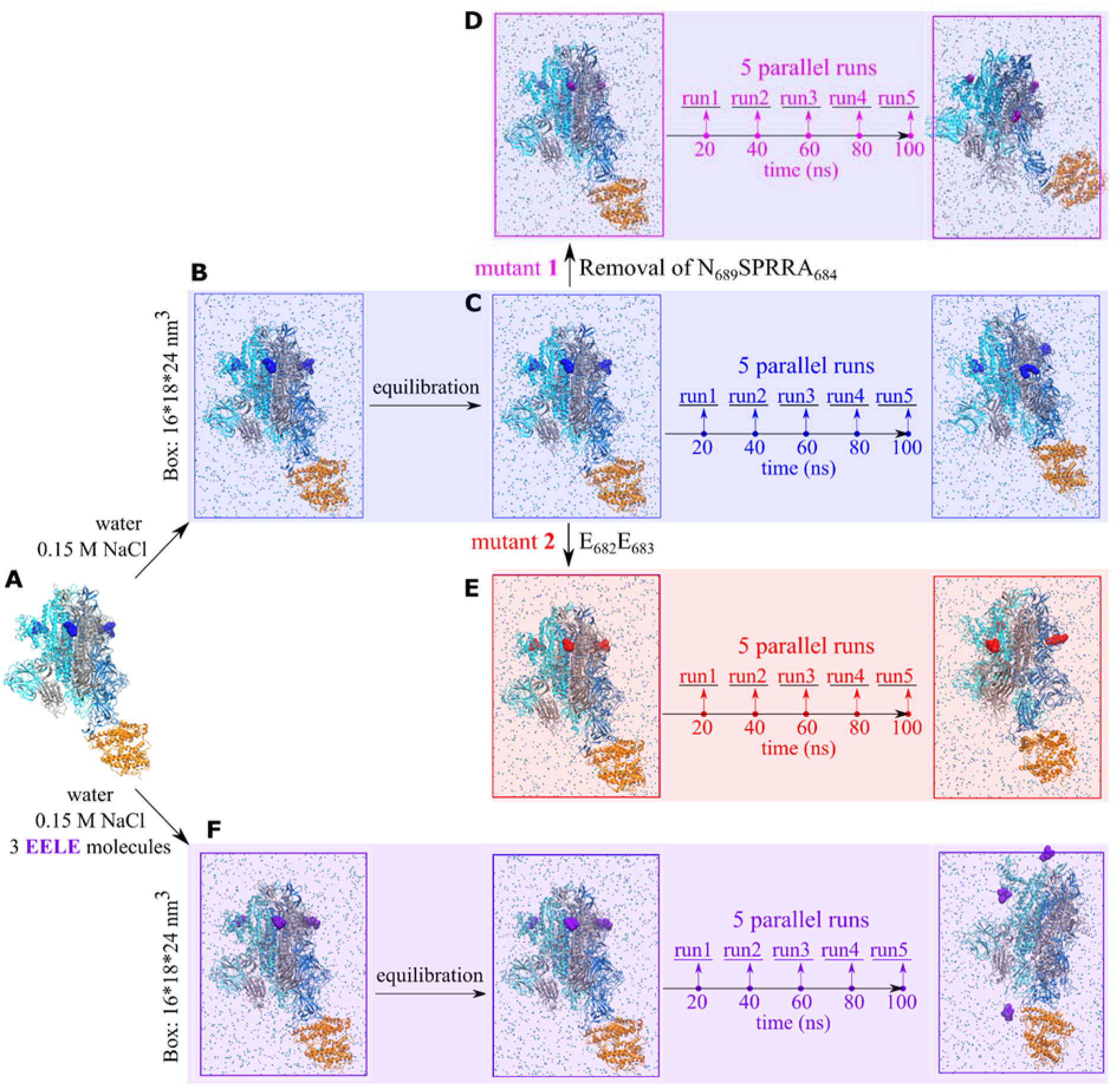
Simulation method in the absence (B-E) and presence (F) of the tetrapeptides EELE. The spike protein-ACE2 complex (**A**) was solvated in a simulation box with 0.15 M NaCl (**B**). The spike protein trimer is colored cyan, silver, and blue, respectively, and ACE2 in orange. Na^+^ and Cl^−^ ions are displayed using small blue and cyan beads, respectively, with water omitted for display. The polybasic cleavage sites (R682RAR_685_) are highlighted with bigger blue beads. The system was equilibrated (**C**). The WT spike protein was mutated by removing N_679_SPRRA_684_ (mutant **1** with R_685_ in purple; **D**) and changing R_682_R_683_ to E_682_E_683_ (mutant **2** with E682EAR_685_ in red; **E**). Each simulation lasted 100 ns (**C**, **D**, **E**). Five parallel runs (10 ns each) were performed based on the configurations at 20, 40, 60, 80, and 100 ns. Same process was applied to the system with EELE tetrapeptides (**F**).

In the production simulation (**Figure 4C**), the periodic boundary conditions were employed in all three dimensions. The neighbor searching was conducted up to 12 Å using the Verlet particle-based method and was updated every 20-time steps. The Lennard-Jones (LJ) 12-6 interactions were switched off from 10 to 12 Å via the potential-switch method in GROMACS. The short-range Coulomb interactions were truncated at the cut-off distance of 12 Å, and the long-range interactions were calculated using the Smooth Particle Mesh Ewald (PME) algorithm ^36, 37^. The temperatures of proteins, ions, and water were separately coupled using the Nosé-Hover algorithm (reference temperature 298K, characteristic time 1 ps). The isotropic Parrinello-Rahman barostat was employed with the reference pressure of 1 bar, the characteristic time 4 ps, and the compressibility 4.5×10^−5^ bar^−1^. All the covalent bonds were constrained, which supported an integration time step of 2.5 fs. These parameters were recommended for the accurate reproduction of the original CHARMM simulation on lipid membrane system ^38^ and were verified in simulations on proteins ^20, 39, 40^ and lipid membranes ^41^. The production simulation lasted 100 ns. Calculation of the root-mean-square deviations of the RBD and ACE2 backbone atoms (**Fig S1** in the Supporting Information) supported that the structures of the RBD and ACE2 were both preserved except that mutant **1** displayed a relatively higher fluctuation. Based on the configurations at 20, 40, 60, 80, and 100 ns, 5 parallel simulations were performed, each of which run 10 ns.

The system had around 700,000 atoms and was computationally expensive. The performance was around 14 ns/day with 8 high-performance computing nodes, each with 24 CPU cores (Intel Haswell E5-2680, 2.5 GHz, 2 × 9.6 GT/s Intel^®^ QPI, 2500 MHz).

The short-range Coulomb and LJ interaction energies (**Table S2** in the Supporting Information) were calculated using the GROMACS program *gmx energy*. The intermolecular hydrogen bonds were calculated using the GROMACS program *gmx hbond*. The structural criteria of hydrogen bond were applied that the donor (D) – acceptor (A) distance *r_DA_* ≤ 3.5 Å and the hydrogen–donor–acceptor angle *θHDA* ≤ 30°.^42, 43^

### Mutants 1 and 2 of SARS-CoV-2 Spike Protein

Based on the equilibrated structure, the wild type spike protein was mutated. For Mutant **1**, the N_679_SPRRA_684_ residues were deleted from all the three subunits of the spike protein trimer. This was to mimic the experimentally reported mutant by Walls et al. ^2^ that the residues of T_678_NSPRRAR_685_ in the wild type spike protein of SARS-CoV-2 were mutated to T_678_--IL--R_685_. Accordingly, six Cl^−^ ions, which were distributed next to the six arginine residues of R_682_R_683_ were removed to neutralize the system. The system was further equilibrated to relax the mutated region, which was followed by the production simulations. The simulation parameters were the same as those in the simulation on the wild type spike protein. Similarly, five parallel simulations were carried out based on the production simulation of 100 ns (**Figure 4D**).

For Mutant **2**, the R_682_R_683_ residues were changed to E_682_E_683_ for all the three subunits of the spike protein trimer. Accordingly, the six Cl^−^ ions, which were distributed next to the six arginine residues (R_682_R_683_) were replaced with Na^+^ ions to neutralize the system. Again, the simulation parameters were the same as those in the simulation on the wild type spike protein. The production simulation lasted 100 ns, where five parallel simulations were performed (**Figure 4E**).

### SARS-CoV-2 Spike Protein-ACE2 Complex in the Presence of Tetrapeptide EELE.

Three tetrapeptides GluGluLeuGlu (EELE) were included, which were initially put next to the three polybasic cleavage sites (R682RAR_685_) of the trimeric wild type spike protein (**Figure 4F**). The spike protein-ACE2 complex and the tetrapeptide EELE were then solved in a simulation box with an edge length of 16×18×24 nm^3^. 0.15 M NaCl was added along with 9 Na^+^ counterions of EELE and 49 Na^+^ counterions of spike protein and ACE2 (**Table S1** in the Supporting Information). The equilibration and productions were the same as whose for the wild type spike protein in the absence of the tetrapeptide. The calculated RBD-ACE2 intermolecular Coulomb and Lennard-Jones interactions are presented in **Table S2** in the Supporting Information.

## Supporting information

Supporting figure of root-mean-square deviations of the RBD and ACE2 backbone atoms. Supporting tables of system compositions and detailed interaction energies.

## ASSOCIATED CONTENT

### Supporting Information

The following files are available free of charge at http://pubs.acs.org/doi/XXX.

## AUTHOR INFORMATION

### Author Contributions

B.Q. developed, performed, and analyzed the simulations. B.Q. and M.O.d.l.C. designed the research, analyzed data, and wrote the manuscript.

### Notes

The authors declare no competing financial interest.

## ACKNOWLEDGMENTS

This work was supported by the U.S. Department of Energy (DOE), Office of Science, Office of Basic Energy Sciences, under Award No. DE-FG02-08ER46539, the Sherman Fairchild Foundation, and the Center for Computation and Theory of Soft Materials at Northwestern University.

