## Supporting figure of root-mean-square deviations of the RBD and ACE2 backbone atoms. Supporting tables of system compositions and detailed interaction energies. for "The Distal Polybasic Cleavage Sites of SARS-CoV-2 Spike Protein Enhance Spike Protein-ACE2 Binding"

### S.I. Supporting Figures

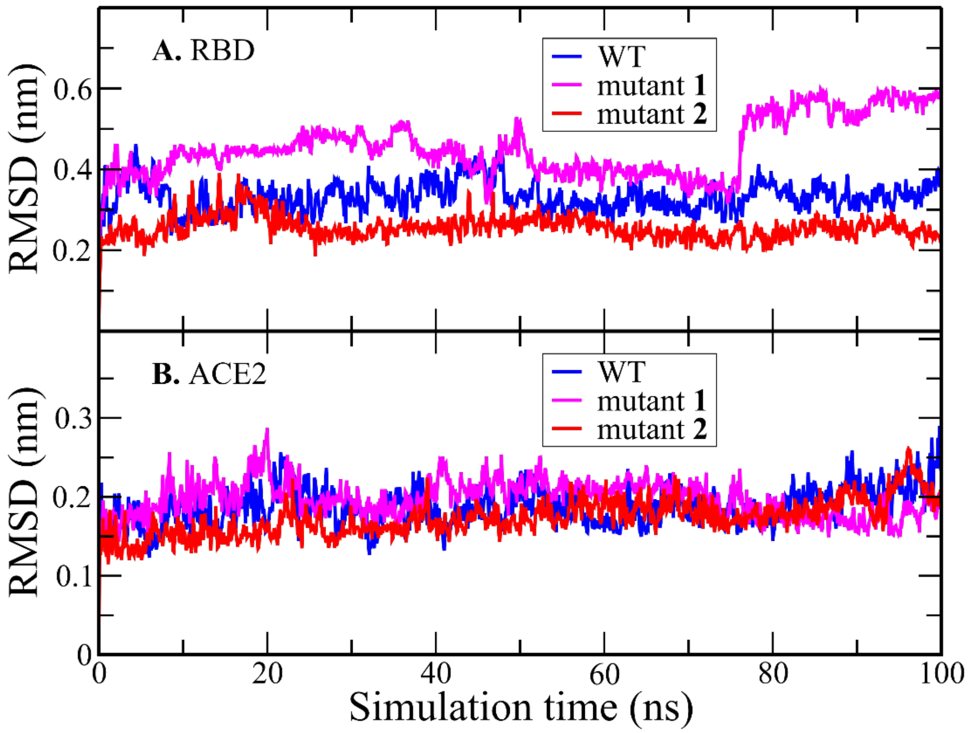


**Figure S1. Room-mean-square deviation (RMSD) of the backbone atoms of (A) the SARS-CoV-2 RBD and (B) the ACE2 receptor.** The RBD structure was similarly preserved except that mutant **1** displayed a relatively higher fluctuation. The ACE2 structure was highly perserved in all three systems.

### S.II. Supporting Tables

**Table S1.** Compositions of the Simulation Systems.

| | #Spike-ACE2 | $\#Na^{+\ e}$ | $\#Cl^{-\ e}$ | $\#H_2O$ | #EELE | Time (ns) [f] |
|---|---|---|---|---|---|---|
| Wild type [a] | 1 | 673 | 624 | 207542 | - | 100 + 5×10 |
| Mutant **1** [b] | 1 | 673 | 618 | 207542 | - | 100 + 5×10 |
| Mutant **2** [c] | 1 | 679 | 618 | 207542 | - | 100 + 5×10 |
| **EELE** [d] | 1 | 682 | 624 | 207465 | 3 | 100 + 5×10 |

a) Wild type spike protein of SARS-Cov-2, where the net charge of each subunit of the trimeric spike protein is -7*e*; and -28*e* for ACE2.
b) Mutant **1** that $N_{679}SPRRA_{684}$ residues were deleted from all three subunits of SARS-CoV-2 spike protein trimer (**Figure 1**). The net charge of each subunit of mutant **1** is -9*e*.
c) Mutant **2** by replacing $R_{682}R_{683}$ with $E_{682}E_{683}$ (**Figure 1**). The net charge of each subunit of mutant **2** is -11*e*.
d) **EELE** = GluGluLeuGlu.
e) 0.15 M NaCl and the counterions of the proteins (wild type spike protein, mutant **1**, mutant **2** and, ACE2) and tetrapeptides.
f) 100 ns simulation and 5 parallel simulations of 10 ns each. (**Figure 4**)

**Table S2.** Short-range Interactions (kJ/mol) for the Wild Type Spike Protein and the Mutants. [a]

| | | **RBD-ACE2** | **Spike-ACE2** | **Spike-EELE** | **Spike-$Na^+$** | **Spike-$Cl^-$** | **Spike-$H_2O$** | **ACE2-$Na^+$** | **ACE2-$Cl^-$** | **ACE2-$H_2O$** |
|---|---|---|---|---|---|---|---|---|---|---|
| Wild type [b] | Coul. | -490 ± 70 | -570 ± 70 | - | -2200 ± 300 | -2100 ± 300 | -310000 ± 1000 | -590 ± 60 | -120 ± 20 | -51300 ± 400 |
| | LJ | -245 ± 7 | -259 ± 7 | - | 80 ± 20 | 30 ± 10 | -34600 ± 500 | 16 ± 4 | -2 ± 1 | -4060 ± 20 |
| | **SUM** | **-740 ± 70** | -830 ± 70 | - | -2100 ± 200 | -2100 ± 200 | -345000 ± 1000 | -570 ± 60 | -120 ± 20 | -55300 ± 400 |
| Mutant **1** [c] | Coul. | -180 ± | -280 ± 40 | - | -2400 ± 200 | -1700 ± 50 | -310000 ± 1000 | -640 ± 60 | -110 ± 20 | -50900 ± 200 |
| | LJ | -290 ± 20 | -310 ± 20 | - | 90 ± 20 | 3 ± 4 | -35100 ± 400 | 21 ± 9 | -2 ± 1 | -4010 ± 60 |
| | **SUM** | **-470 ± 50** | -590 ± 50 | - | -2300 ± 200 | -1700 ± 60 | -344900 ± 800 | -620 ± 60 | -110 ± 20 | -54900 ± 200 |
| Mutant **2** [d] | Coul. | -330 ± 50 | -350 ± 40 | - | -2200 ± 200 | -1600 ± 100 | -317000 ± 1000 | -800 ± 100 | -130 ± 30 | -52000 ± 500 |
| | LJ | -260 ± 4 | -284 ± 6 | - | 68 ± 9 | 6 ± 6 | -34700 ± 200 | 27 ± 8 | -1 ± 2 | -4130 ± 40 |
| | **SUM** | **-590 ± 40** | -630 ± 30 | - | -2200 ± 200 | -1600 ± 100 | -352000 ± 1000 | -700 ± 100 | -130 ± 30 | -56100 ± 400 |
| **Wild type + EELE** [e] | Coul. | -240 ± 50 | -260 ± 40 | -200 ± 200 | -2100 ± 200 | -1600 ± 200 | -312000 ± 2000 | -580 ± 80 | -130 ± 20 | -52400 ± 500 |
| | LJ | -250 ± 20 | -260 ± 20 | -70 ± 60 | 70 ± 20 | 5 ± 7 | -34600 ± 300 | 20 ± 7 | 0 ± 2 | -4080 ± 30 |
| | **SUM** | **-490 ± 50** | -520 ± 50 | -300 ± 300 | -2000 ± 200 | -1600 ± 200 | -346000 ± 2000 | -560 ± 70 | -130 ± 20 | -56500 ± 400 |

a) Interactions are calculated up to 1.2 nm, which is the cut-off distance for short-range Coulomb (Coul.) and LJ 12-6 interactions in the simulations. Error bars stand for the standard deviations from five parallel runs (**Figure 4**).
b) Wild type spike protein of SARS-CoV-2.
c) Mutant **1** that $N_{679}SPRRA_{684}$ residues are deleted from SARS-CoV-2 spike protein (**Figure 1**).
d) Mutant **2** by replacing $R_{682}R_{683}$ with $E_{682}E_{683}$ (**Figure 1**).
e) Wild type spike protein in the presence of tetrapeptide **EELE**.
